# Differential impact of brain network efficiency on post-stroke motor and attentional deficits

**DOI:** 10.1101/2022.05.23.493043

**Authors:** Giorgia G. Evangelista, Philip Egger, Julia Brügger, Elena Beanato, Philipp J. Koch, Martino Ceroni, Lisa Fleury, Andéol Cadic-Melchior, Nathalie Meyer, Diego de León Rodríguez, Gabriel Girard, Bertrand Léger, Jean-Luc Turlan, Andreas Mühl, Philippe Vuadens, Jan Adolphsen, Caroline Jagella, Christophe Constantin, Vincent Alvarez, Joseph-André Ghika, Diego San Millán, Christophe Bonvin, Takuya Morishita, Maximilian J. Wessel, Dimitri Van de Ville, Friedhelm C. Hummel

## Abstract

**Background:** Most studies on stroke have been designed to examine one deficit in isolation, yet survivors often have multiple deficits in different domains. While the mechanisms underlying multiple-domain deficits remain poorly understood, network-theoretical methods may open new avenues of understanding.

**Methods:** 50 subacute stroke patients (7±3days post-stroke) underwent diffusion-weighted magnetic resonance imaging and a battery of clinical tests of motor and cognitive functions. We defined indices of impairment in strength, dexterity, and attention. We also computed imaging-based probabilistic tractography and whole brain connectomes. Overlaying individual lesion masks onto the tractograms enabled us to split the connectomes into their affected and unaffected parts and associate them to impairment.

**Results:** To efficiently integrate inputs from different sources, brain networks rely on a “rich-club” of a few hub nodes. Lesions harm efficiency, particularly when they target the rich-club. We computed efficiency of the unaffected connectome, and found it was more strongly correlated to impairment in strength, dexterity and attention than efficiency of the total connectome. The magnitude of the correlation between efficiency and impairment followed the order attention > dexterity ≈ strength. Network weights associated with the rich-club were more strongly correlated to efficiency than non-rich-club weights.

**Conclusions:** Attentional impairment is more sensitive to disruption of coordinated network activity between brain regions than motor impairment, which is sensitive to disruption of localized network activity. Providing more accurate reflections of actually functioning parts of the network enables the incorporation of information about the impact of brain lesions on connectomics contributing to a better understanding of underlying stroke mechanisms.

## Introduction

It has long been acknowledged that different brain regions are linked together in complex patterns, making networks a natural mathematical model for the brain, with regions of the brain serving as nodes and edges being weighted according to structural characteristics.^1–4^ Furthermore, neuroimaging evidence in humans suggests that stroke is a network disease, indicating that using network theory as the basis of a model for stroke^5–7^ might significantly enhance the understanding of stroke, its deficits and the recovery therefrom.

It is useful to think of networks on a spectrum between regularity and randomness. Connectivity in regular networks tends to feature well-defined local communities, while in random networks it tends to feature one global community with costly long-distance connections;^8^ brain networks occupy the zone on the spectrum in which the tradeoff between global integration and local segregation is optimal.^1^ They optimize the tradeoff with an architecture featuring a small set of hub nodes called a “rich-club” (RC).^1,9^ These hubs are “rich” because they are strongly connected to nearby nodes, and form a “club” because they are strongly connected to each other. The RC can be thought of as a backbone for global connectivity, and therefore “attacks” (e.g., stroke lesions) against it will have a greater impact on global connectivity than random attacks of similar magnitude.^9^

When brain networks are “attacked” by a stroke, the effect on global integration can be considerable, particularly when the attack focuses on the RC.^10^ Likewise, the effect on behavioral function can be significant, particularly on cognitive functions, such as attention, that rely heavily on global integration as opposed to those functions whose neurological substrate is more localized, such as sensorimotor functions.^11^

Structural connectomics relies on models of white matter (WM) tractography computed from diffusion-weighted imaging (DWI). In areas with high axon density, water molecule diffusion has a strong preference for the direction of the axons, i.e., high anisotropy. By chaining together high-anisotropy voxels in the relevant directions, one can extract a model for WM tracts. Typically, stroke-lesioned brain tissue undergoes significant changes including liquefactive necrosis, reducing the anisotropy with consequences on the modelled WM tracts. Nonetheless, many paths pass through lesioned tissue, even though they might not correspond to a functioning axon bundle.

We set out to answer two questions. First, is the understanding of connectomics significantly enhanced by the lesion structure information derived from DWI? To answer this question, we defined the structural connectome with and without explicit lesion information, respectively, and compared the data to behavioral metrics.

Second, how strongly are network-theoretic notions of global connectivity or RC integrity associated with stroke-induced impairment in different behavioral domains? We hypothesized that global connectivity and RC integrity will be associated with these behavioral functions, but that this association will be stronger in the “less-localized” attentional domain than in the “more-localized” motor domain. To test this hypothesis, we defined indices of impairment in motor and attentional functioning, and correlated them with a mathematically-defined notion of global efficiency (GE).

## Methods

### Patients

We recruited eighty-five stroke patients admitted between 2018 and 2021 to the stroke unit of the Hospital of Valais in Sion, Switzerland, of which ***N* = 50** completed both imaging sessions and behavioral tests and were therefore included in the study (Fig 1B). The inclusion criteria included being older than eighteen years, presence of a motor deficit, and absence of contraindications for MRI or noninvasive brain stimulation (NIBS). Exclusion criteria included requests not to be informed in case of incidental findings, inability to provide informed consent, severe neuropsychiatric or medical disease, history of seizures, pregnancy, regular use of narcotic drugs, presence of implanted devices incompatible with MRI or transcranial magnetic stimulation, use of medication that interacts with NIBS, severe sensory, musculoskeletal or cognitive deficit incompatible with understanding instructions or performing experiments. For detailed patient characteristics please see Table 1. The lesion locations were representative of the overall stroke patient population as shown in the lesion heatmap (Fig 1A), and were not used as a selection criterion. All patients gave written informed consent at the time of enrolment. The current data was acquired in the framework of a larger project (TiMeS project work package 1) and all research was approved by the local ethical committee swissethics (approval number 2018-01355).

**Figure 1:**
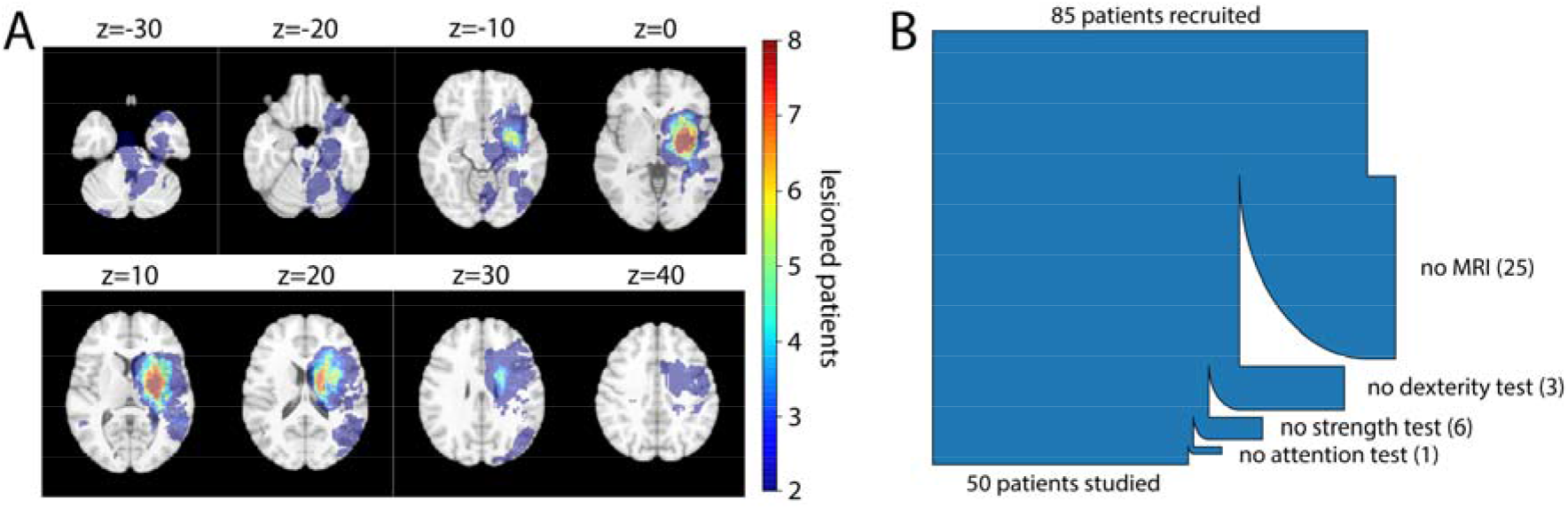
Cohort characteristics. (A) Lesion heatmap where all patients’ lesions are co-registered to the MNI template brain and flipped onto the right hemisphere. Titles refer to z coordinates in the MNI space, i. e. mm superior to the anterior commissure. (B) Patient flowchart. Please note that of the recruited patients, only those who underwent MRI and all behavioral tests were included in the study.

|  | Value<sup>a</sup> | Unit |
| --- | --- | --- |
| Sex | 38 male / 12 female | Patients |
| Age | 65.2 ± 13.7 | Years |
| Time of MRI | 4 ± 2 | Days post-stroke |
| Time of behavioral tests | 7 ± 3 | Days post-stroke |
| Paretic side | 28 left / 22 right | Patients |
| Thrombolysis | 17 yes / 33 no | Patients |
| NIHSS | 5.8 ± 5 | Points |
| FMUE | 53.6 ± 18.1 | Points |
| Fist grip strength | 28.1 ± 15.8 | Kg force |
| Pinch grip strength | 3.6 ± 2.2 | Kg force |
| Key grip strength | 5.8 ± 3.3 | Kg force |
| Box & Block test | 34.6 ± 18.6 | Blocks |
| Purdue pegboard test | 6.7 ± 4.4 | Pegs |
| CTT part A | 77 ± 56 | Seconds |
| CTT part B | 143 ± 78 | Seconds |
| RT on alertness test of TAP | 342 ± 130 | Milliseconds |
| RT on divided attention test (single condition) of TAP | 1012 ± 205 | Milliseconds |

<sup>a</sup>Categorical values are given as breakdowns by level of the category, separated by forward slashes. Numeric values are given with mean and standard deviation, separated by ±. NIHSS = National Institutes of Health Stroke Scale.
FMUE = Fugl-Meyer Upper Extremity score
RT = Reaction Time

### Clinical Assessment

Each patient underwent MRI at the subacute stage in addition to a battery of clinical tests of motor and cognitive function. The focus of this work was on motor and attentional functions. Motor strength was measured by performing the fist, grip and pinch strength test^12^ on both hands, motor dexterity by performing the Box&Block^13^ and Purdue^14^ tests. Attentional functions were measured using the Test of Attentional Performance (TAP),^15^ the Color Trail Test (CTT) parts A and B,^16,17^ and the Bells test.^18^ These tests were selected for fitting in the Sohlberg-Mateer model, which involves the use of five types of attention of increasing difficulty.^19^

### MRI Data Acquisition

All images were acquired using a 3T MAGNETOM Prisma (Siemens Healthcare, Erlangen, Germany) with a 64-channel head and neck coil.

T1-weighted anatomic images were acquired using 3D magnetization-prepared, rapid acquisition gradient-echo sequence (MPRAGE) with the following parameters: 192 axial slices, response time = 2300ms, echo time = 2.96ms, flip angle = 9°, voxel size = 1×1×1mm, field of view = 256×256mm.

For the DWI, diffusion gradients with five different gradient strengths (b-values = [300,700,1000,2000,3000]s/mm^2^; shell-samples = [3,7,16,29,46]) were obtained in 101 non-collinear directions distributed equally over the brain in 84 axial slices. The images were acquired using the pulsed gradient spin echo technique with the following parameters: repetition time = 5000ms, echo time = 77ms, field of view = 234×234mm, voxel resolution=1.6×1.6×1.6mm, readout bandwidth = 1630Hz/pixel, GRAPPA acceleration factor = 3.

Seven T2-weighted images without diffusion weighting (b=0s/mm^2^) were acquired, including one in opposite phase encoded direction.

### Lesion Segmentation

All the lesion masks were hand-drawn using MRview from MRtrix3^20^ and subsequently verified by a neurologist. This enabled us to also compute, for each streamline, the binary value indicating whether or not the streamline passed through the lesion as described in previous work.^10^

### Image Analysis

Tissue partial volume maps were estimates from the T1-weighted image and registered to the average b0 image using FSL.^21^ FreeSurfer was used to obtain a brain parcellation including 74 cortical areas per hemisphere (Destrieux atlas), subcortical areas (thalamus, caudate, putamen, hippocampus, amygdala), the cerebellum, and a subdivision of the brainstem (midbrain, pons, medulla), yielding 163 brain areas.^22^ The voxels corresponding to the lesion were stamped out and replaced by the mirrored voxels of the contralateral side.

The DWI were preprocessed using MRtrix3,^20^ and FSL^21^(Gibbs ringing, motion, field inhomogeneity, susceptibility-induced off-resonance field, eddy currents and bias-field correction). Multi-shell multi-tissue constrained spherical deconvolution^23^ was used to estimate the fiber orientation distributions within each voxel. Whole-brain probabilistic tractography was performed using the MRtrix3 second-order integration over fiber orientation distribution method,^20^ initiating streamlines in all voxels of the WM. Streamline tracking parameters were set to default values, except the minimum streamline length of 1.6mm. For each dataset, 1 million streamlines were selected with both endpoints in the individual cortical or subcortical mask using the Dipy software package.^24^ Every streamline was weighted fitting the underlying diffusion compartment model using a Stick-Ball-Zeppelin^25^ model using COMMIT, a practice designed to boost the anatomical accuracy of the tractography mainly by avoiding or down-weighting false positives.^26^

### Total and Unaffected Connectome

For each patient, a structural connectome was built with 13,202 pairs of areas obtained through the parcellation. ^10,27^ As in our previous work, we did not limit the analyses to considering the numbers of streamlines between areas, a method which suffers from serious pitfalls.^28^ Instead, we computed whole brain connectomes as follows: for each pair of regions of interest, we compute the sum of the COMMIT weights of the streamlines that run between the two regions. Taken together, the COMMIT weighting and the large number of streamlines ensure that the estimated diffusion connectivity stabilizes, mitigating the pitfalls by improving the robustness and reproducibility of diffusion connectivity estimations.^26,29^

Then we use the binary map indicating whether or not the streamline passed through the lesion to split the full connectome *C_total_* into the sum

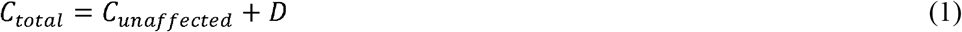

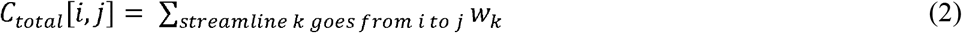

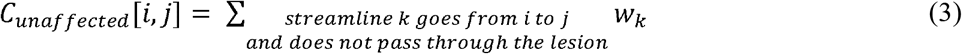

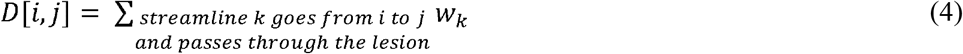

In this way, we are able to incorporate the lesion more explicitly and by only considering streamlines which are not lesioned, simultaneously encode true structural connectivity and mitigate the potential for artifacts of e.g., Wallerian degeneration.^30^

### Rich-club, Edge Weight, and Node Weight

We considered the same set of nodes as found by van den Heuvel and Sporns to form the RC, namely the bilateral precuneus, superior frontal cortex, superior parietal cortex, hippocampus, putamen, and thalamus.^9^ Finally, we split the edges into three groups (Fig 2): pure RC connections (between RC nodes); feeder connections (between a RC and a non-RC node); and local connections (between non-RC nodes).

**Figure 2:**
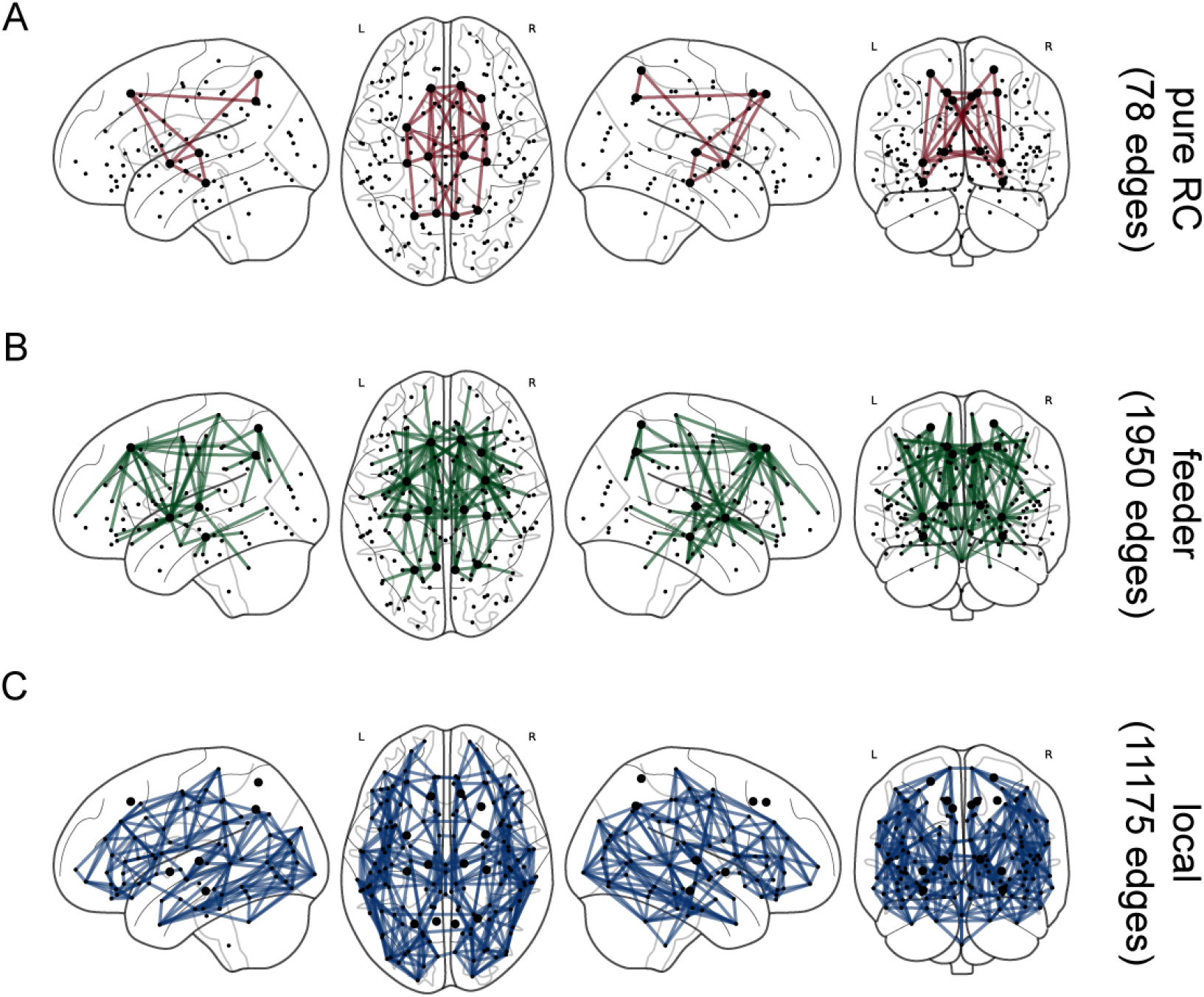
Connectome nodes and edges on glass brain. Our parcellation contains 163 nodes, depicted as dots on the brain. 18 of these nodes, shown as enlarged dots, are the RC nodes. The edges are partitioned into pure RC (**A**, red), feeder (**B**, green), and local (**C**, blue) types.

Each edge has a weight; each node has a weight, defined to be the sum of the weights of all edges linked to that node.

### Dimensionality Reduction

The behavioral metrics of strength, dexterity and attention all were represented by multiple features, which implies a need for dimensionality reduction. Popular methods include principal component analysis and nonnegative matrix factorization (NMF);^31^ because our data are nonnegative by nature and distance from zero impairment has a relevant meaning, we chose NMF. The amount of information that is lost by reducing features is captured by calculating the proportion of variance accounted for (VAF) by the low-dimensional representation.^31^

### Behavioral Metrics

To measure strength impairment, we consider the average force in three trials exerted by the patients on both hands, and calculate a normalized impairment metric as follows:

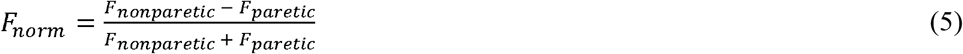

in a fist grip, a pinch grip and a key grip,^12^ giving the three features *FIST_norm_,PINCH_norm_*, and *KEY_norm_*. These features are bounded between −1 and 1, where −1 means only the paretic hand exerts force, 0 means both hands exert equal force, and 1 means only the nonparetic hand exerts force. While mildly negative values are possible, they are unlikely to be clinically meaningful, so we set negative values to zero in order for the features to be nonnegative. Patients missing all three of *FIST_norm_,PINCH_norm_* and *KEY_norm_* were excluded; those who were missing some but not all were replaced by the mean of all non-missing data in the respective field. Finally, we used NMF (*VAF* = 97%) to reduce the three features into one strength impairment index

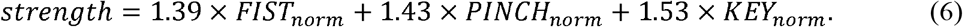

To measure dexterity impairment, we consider the number of fine motor tasks performed by the patient with both hands in a given time limit. As before, we calculate the normalized functional metric

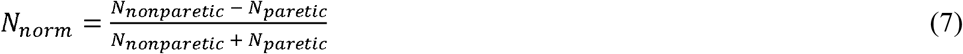

in the box-and-block test^32^ and the Purdue pegboard test,^14^ set negative values to zero, impute missing data, and use NMF (*VAF* = 97%) to reduce the two features into one dexterity impairment index

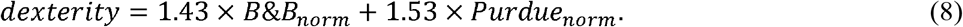

To measure functional deficits in attention, we began with the widely-used model of Sohlberg-Mateer,^19^ which suggests measuring five tasks of increasing difficulty, as described in Table 2:

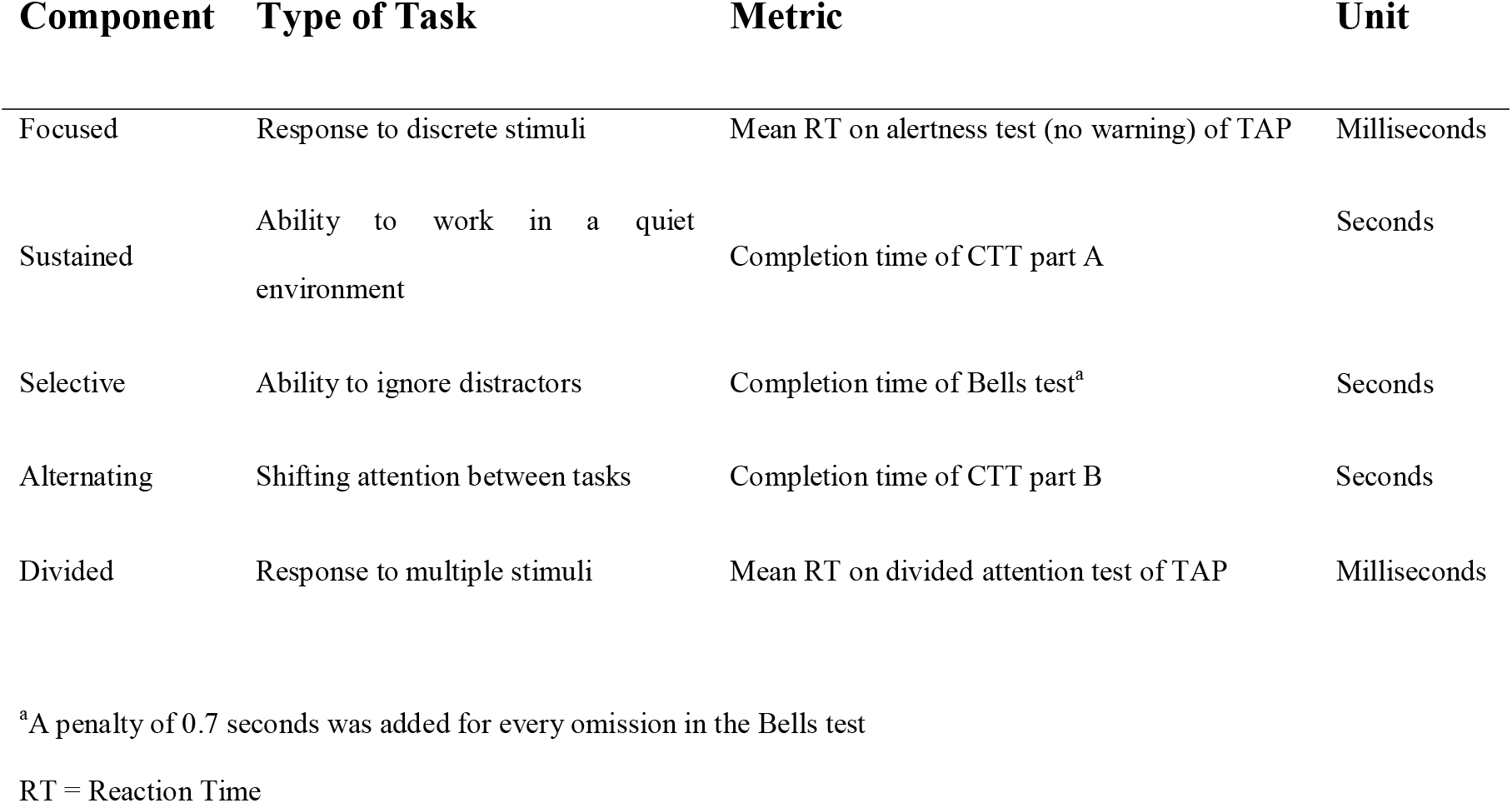

| Component | Type of Task | Metric | Unit |
| --- | --- | --- | --- |
| Focused | Response to discrete stimuli | Mean RT on alertness test (no warning) of TAP | Milliseconds |
| Sustained | Ability to work in a quiet environment | Completion time of CTT part A | Seconds |
| Selective | Ability to ignore distractors | Completion time of Bells test<sup>a</sup> | Seconds |
| Alternating | Shifting attention between tasks | Completion time of CTT part B | Seconds |
| Divided | Response to multiple stimuli | Mean RT on divided attention test of TAP | Milliseconds |

<sup>a</sup>A penalty of 0.7 seconds was added for every omission in the Bells test
RT = Reaction Time

We normalize each of these features to have maximal value 1, impute missing values, and use NMF (*VAF* = 94%) to reduce the five normalized features to one attention impairment index

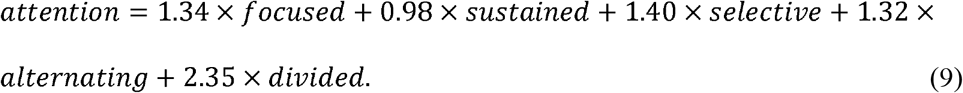

### Global Efficiency

Most networks lie on a spectrum between local segregation and global integration. Many biological networks are “small-world” networks, which have both the clustered nature of locally segregated networks and the short path length of globally integrated networks.^33,34^ The GE of a network with *n* nodes is defined by Rubinov and Sporns^34^ as

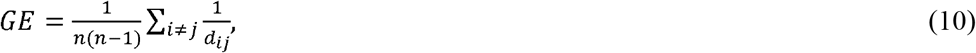

where *d_ij_* is the length of the shortest path between nodes *i* and *j*, ie. the smallest sum of reciprocal edge weights in any path from *i* to *j*. GE of the brain networks were computed using MATLAB’s Brain Connectivity Toolbox.^34^

### Data availability

Data will be made available upon reasonable request.

## Results

We computed the weighted GE of each patient’s total connectome *C_total_* and unaffected connectome *C_unaffected_*; and then correlated these with strength, dexterity and attention impairment.

As GE was correlated with lesion volume, we considered lesion volume as a confounder. To establish the wisdom of using GE nonetheless, we performed recursive feature elimination on linear regression models of type *impairment ~ lesion volume* + *GE_total_* + *GE_unaffected_* and in all three domains, the last predictor variable remaining was *GE_unaffected_*.

All correlations were estimated using bootstrap resampling^35^ with 10,000 iterations, i.e., at each iteration we drew 50 patients from our sample of 50 (with replacement) and computed the respective correlations on the subsample. We computed the Pearson correlation between *GE_total_* and impairment (Fig 3, blue; strength: *r* = −.20, *P* = .07, dexterity: *r* = −.11, *P* = .25, attention: *r* = −.41, *P* = .0001); and between *GE_unaffected_* and impairment (Fig 3, green; strength: r = −.30, *P* = .02, dexterity: *r* = −.30, *P* = .05, attention: *r* = −.55, *P* < 0.001). The negative correlations are expected, given that GE is known to contribute to better functional outcomes;^11,36^ p-values are proportions of bootstrap correlations that were nonnegative.

**Figure 3:**
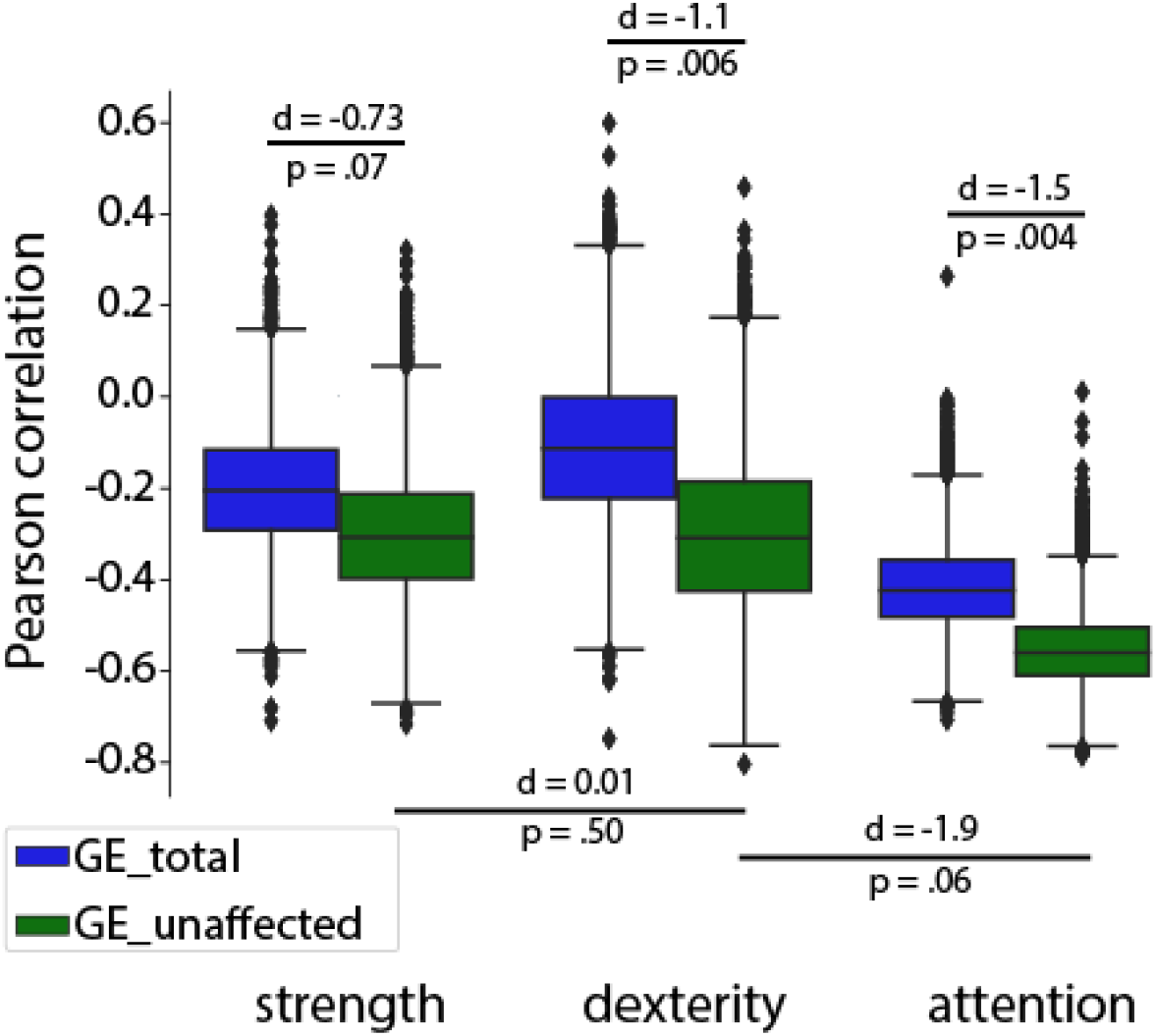
Bootstrapped correlations between connectome GE and impairment. Boxplots show bootstrapped Pearson correlations on the ordinata between GE and the three types of impairment on the abscissa. Blue plots refer to GE_total_, while green plots refer to GE_unaffected_. Note that all correlations tend to be negative and that correlations with GE_unaffected_ tend to be stronger (i.e. more negative) than those with GE_total_. Note also that GE tends to be more strongly correlated to attention impairment than to strength or dexterity impairment, while it seems to be equally strongly correlated to strength and dexterity impairment.

We computed effect sizes for the difference between *r*(*GE_unaffected_, impairment*) and *r*(*GE_total, impairment_*) across bootstrap iterations using Cohen’s *d* statistic rather than Student’s t due to the latter showing inflated effect sizes with large datasets;^37^ p-values were computed by probability of superiority.^38^ *GE_unaffected_* was more strongly correlated with impairment than *GE_total_* (strength: *d* = −0.73, *P* = .07, dexterity: *d* = −1.1, *P* = .006, attention: *d* = −1.5, *P* = .004). We did not control for age, as age was found not to be a meaningful covariate in previous studies of older healthy subjects.^11^

It is known from work of van den Heuvel and Sporns^9^ that attacks on pure RC edges result in more loss of GE than attacks of similar magnitude on feeder or local edges. Conversely, we expected that greater integrity of pure RC edges should correspond to greater GE in the network, and this is the case (Fig 4A, left: one-way ANOVA *F* = 82.1, *P* < 0.001). Greater integrity of RC nodes also corresponds to greater GE in the network (Fig 4B, left: two-sample t test *T* = 2.6, *P* = .009).

**Figure 4:**
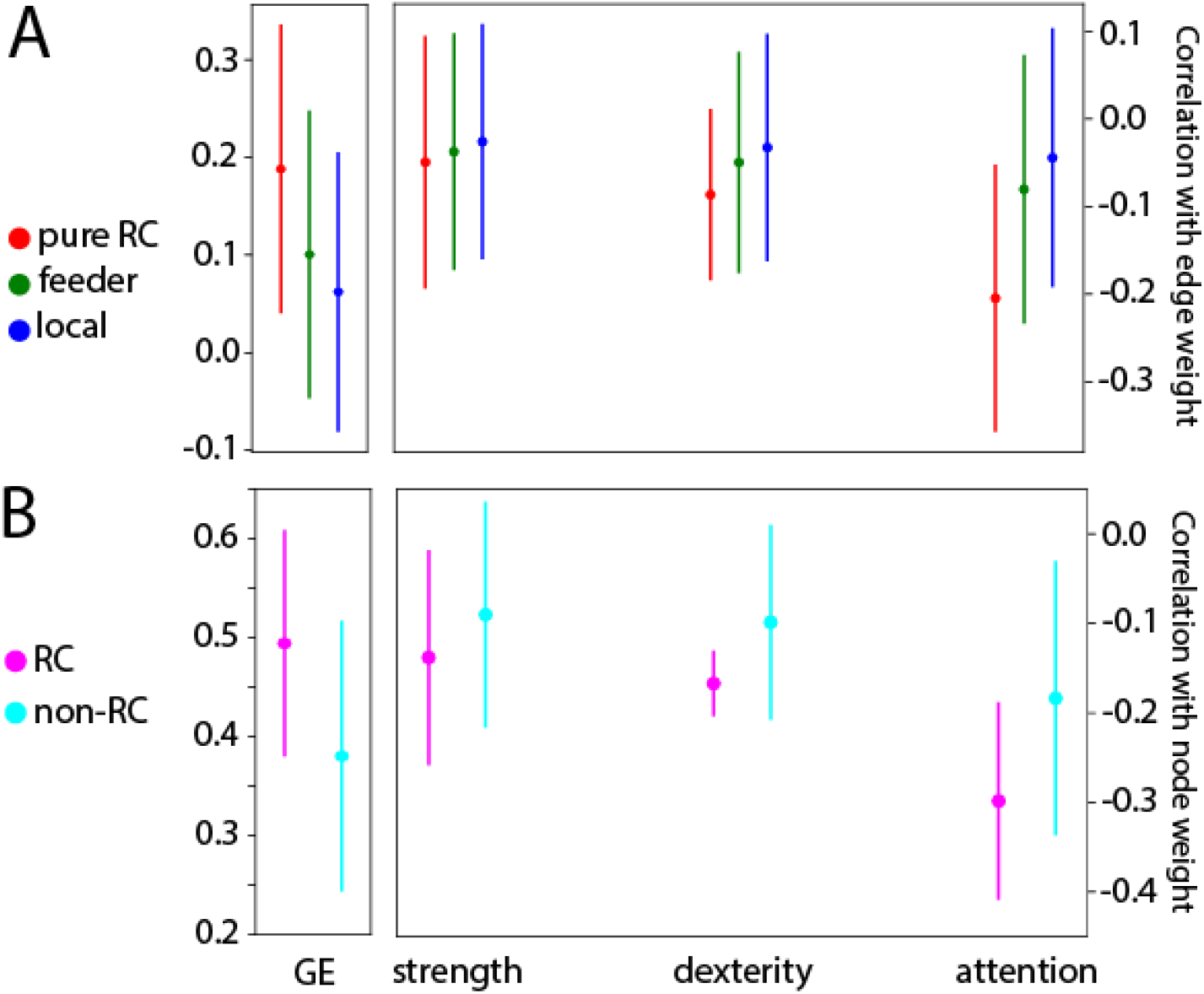
Correlations between connectome GE, graph weights, and impairment. Plots show mean and standard deviation. (**A**, left) Correlations between edge weight and GE_unaffected_. Please note that the order pure RC > feeder > local holds. (**A**, right) Correlations between edge weight and impairment. Please note that the order pure RC < feeder < local holds in all domains, and that the gap is largest in the attentional domain. (**B**, left) Correlations between node weight and GE_unaffected_. Please note that RC nodes have stronger correlation to GE_unaffected_ than non-RC nodes. (**B**, right) Correlations between node weight and impairment. Please note that RC nodes have stronger correlations to behavior than non-RC nodes and that the gap increases between strength, dexterity, and attention.

It has been shown that among healthy older subjects, when considering correlations to attention, which requires integration of inputs from across the brain, pure RC edges are strongest, then feeder, then local. However, no such order exists for correlations to visual processing, which is more localized.^11^ The same holds when considering RC nodes as opposed to non-RC nodes.

We have found analogous results to those reported by Baggio and colleagues^11^ in healthy older adults. Pure RC edge weights have more negative (i.e. stronger) correlations to behavior than feeder and local edges, and this gap is smallest for strength and largest for attention (Fig 4A, right: one-way ANOVA. Strength: *F* = 6.5, *P* < .001; dexterity: *F* = 19.7, *P* < 0.001; attention: *F* = 89.4, *P* < 0.001). Similarly, RC node weights have stronger correlations to behavior than non-RC node weights, and this gap is smallest for strength and largest for attention (Fig 4B, right: two-sample t test. Strength: *T* = 0.7, *P* = .474; dexterity: *T* = 2.0, *P* = .048; attention: *T* = 3.0, *P* = .003).

## Discussion

Given the complex interactions between different brain areas, mathematical tools for complex network analyses offer an exciting opportunity to better understand mechanisms underlying neurological disorders, especially when current findings strongly support the maxim that many neurological disorders, including stroke, are network disorders.^5,39^ Therefore, by evaluating the patient’s specific brain connectivity, connectomics has great potential to yield important insights into post-stroke impairment and recovery mechanisms.

Our data suggest there are considerable differences in the mechanisms underlying deficits in the motor and attentional domains. These differences provide justified optimism that while treatments designed to promote more globally efficient brain networks are likely to have benefits in treating many deficits, this is particularly true of attentional deficits.

While the stroke patients in this study were selected for their motor deficit, many also showed cognitive/attentional deficits (e.g., 39/50 pathological on MOCA). It has been suggested from studies of older healthy subjects^11^ and stroke patients^40^ that cognitive functions, such as attention, memory or language functions, are more heavily reliant on integration of inputs from different parts of the brain than functions such as motor or visual ones, which reside in “more specialized” local brain networks. This important assumption has been confirmed by our findings that *GE_unaffected_* is more strongly correlated to attention than to motor strength or dexterity, highlighting the reliance of the attentional domain on larger-scale emergent dynamics. Note that the correlation between *GE_unaffected_* and strength does not differ significantly from that between *GE_unaffected_* and dexterity, highlighting the fact that the motor domain, whether for pure strength production or for more fine motor skills, is less reliant on emergent (larger-scale) dynamics than attention is. Mathematical modeling conducted by Sporns and van den Heuvel has found that the resilience of brain networks to “attack” varies depending on the place of attack, indicating that attacks on pure RC edges result in greater drops in GE than other (non-RC) attacks of similar magnitude.^9^ Accordingly, we found that RC node weight and pure RC edge weight were more strongly correlated to attention than to motor functions, suggesting that the importance of the RC is derived from its disproportionate impact on GE.

Our approach adds personalization to traditional connectomics by separating fibers according to whether or not they are impacted by the patient’s particular lesion. This approach has borne fruit, as the unaffected connectome is more strongly associated with behavior than the traditional total connectome. From a hypothesis-driven perspective, the importance of including a lesion profile in the measurement of brain networks is obvious as WM tracts can be detected by DWI-derived tractography, though probably with more diffusivity inhomogeneities, even if they are interrupted by a lesion (particularly in the early post-stroke period). Such tractography by itself fails to acknowledge that if tracts are interrupted by the lesion, their ability to transmit information is compromised and they will not contribute to the normal functioning of the network. From a data-driven perspective, it has been found that *GE_total_* does not differ significantly either over time, or even between stroke patients and healthy controls,^40^ casting doubt on the value of *GE_total_* as a biomarker. By contrast, we provide evidence that *GE_unaffected_* differs significantly from *GE_total_*, and that in stroke patients *GE_unaffected_* is more strongly correlated to behavior than *GE_total_* is.

Consequently, we find that explicitly discarding tracts that pass through the lesion as inoperative ensures that the unaffected connectome is a more accurate reflection of true patterns of connectivity than the traditional structural connectome, and thus a better candidate as a stroke-related biomarker.

### Limitations

The primary drawback of our approach is that splitting the total connectome into its affected and unaffected parts requires that one draw lesion masks and overlay them onto the tractography. This imposes considerable additional work, yet we are convinced that the benefits in terms of relevance to behavior and understanding of mechanisms are worth the cost. In addition to being time-consuming and labor intensive, they require substantial anatomical expertise, which might introduce a considerable source of variability; both on an inter-rater and (to a lesser extent) test-retest basis.^41^ While the use of machine learning algorithms to delineate lesions holds some promise, it is a sufficiently difficult task that at the time of writing, human-drawn lesions remain the gold standard, with even the best available algorithms failing to come close to inter-rater levels of agreement with humans.^42^

### Future Work

This study was conducted cross-sectionally, however longitudinal evaluation of parameters of network efficiency in the affected and unaffected parts of the network will open novel opportunities to study the mechanisms underlying recovery of multi-domain (e.g., motor and attention) post-stroke deficits. It will provide novel insights into the reorganization of structural brain networks, how these reorganizational patterns relate to recovery of behavioral functions, and whether they allow prediction of outcome or treatment stratification. In particular, it would be worth investigating whether among our cohort, patients’ increase in GE over time was associated with recovery from their impairment, particularly attentional impairment.

There are biological reasons to expect that this might occur. Reparative axonal sprouting is characterized by growth of long-distance connections,^43^ the precise type of connections that contribute most to GE. It has also been found to be clearly associated with post-stroke behavioral recovery.^43^

## Conclusions

While the patients in our cohort were selected for motor deficits, most of them also had an attention deficit, which can have an additional impact on the recovery process. This considerable overlap between motor and attentional deficits implies the need for finer-grained discriminators between multidomain deficits and their underlying mechanisms.

Here, we suggest structural connectivity analyses of RC properties with a focus on affected and unaffected parts of the network to better characterize these multi-domain deficits in the subacute stage after stroke. These analyses allow the following conclusions: First, current structural connectomics approaches use DWI to trace axon fiber bundles and simply rely on lesion-induced tract inhomogeneities. However, this implicit consideration of the lesion is suboptimal, particularly in the acute and subacute phase as it might lead to the tractography finding tracts that are no longer functional. Therefore, the results support the importance of explicitly overlaying lesion masks onto the tractography to be able to split the structural connectome into its unaffected (well-functioning) and affected (non-functional) parts to determine which parts of the brain network are actually relevant to residual functions and impairment. Secondly, based on this approach, the results suggest that in stroke patients, attention is more sensitive to non-integrity of the connectome (especially the RC) and the resulting deficiency of GE than motor functions, strongly underscoring the differential importance of RC network properties for different behavioral functions. The results further confirm and are consistent with reports in healthy subjects that the neural substrate underlying motor functions is rather localized, while that underlying attention is based on a more global representation in terms of RC organization.

## Non-standard Abbreviations and Acronyms

DWI: Diffusion-Weighted Imaging
WM: White Matter
GE: Global Efficiency
NIBS: Non-Invasive Brain Stimulation
NMF: Nonnegative Matrix Factorization
RC: Rich-Club
TAP: Test of Attentional Performance
CTT: Color Trail Test
VAF: Variance Accounted For

## Acknowledgements

We thank Silvia Avanzi for her excellent support during the recruitment and data acquisition process.

